# A Rapid Anterior Auditory Processing Stream Through the Insulo-Parietal Auditory Field in the Rat

**DOI:** 10.1101/2023.09.12.557409

**Authors:** Maciej M. Jankowski, Mousa Karayanni, Mor Harpaz, Ana Polterovich, Israel Nelken

## Abstract

The insular cortex is believed to be involved in a wide range of auditory functions in the mammalian brain. We studied the organization and basic response properties of auditory neurons in insular cortex and adjacent areas by recording responses to sound stimuli in anesthetized rats. Auditory neurons were present in an insulo-parietal auditory field that spans the boundary between the posterior insula, particularly in the granular insular cortex and the ventral part of the secondary somatosensory cortex. Neurons in this field had narrow tuning, were preferentially tuned to relatively low frequencies (<16 kHz), and had short response latencies. Intriguingly, some auditory units in this insulo-parietal field displayed shorter onset latencies than the minimal latency in primary auditory cortex. At the same time, these units showed weaker sensitivity to deviance than units in primary auditory cortex. These results establish the existence of a rapid information stream through the insulo-parietal cortex that may parallel the pathway through the primary and anterior auditory fields.

## Introduction

The insular cortex has been shown to be involved in a wide range of auditory functions in the human brain. Strokes in insular cortex are associated with hearing deficits, particularly in temporal resolution and sequencing^1^. Uni- and bilateral vascular events that include the insulae may result in auditory agnosia to environmental sounds, speech, and music. Symptoms often resolve to a varying degree over a few months to years^2–5^. In some cases, persistent hyperacusis is observed^6^.

Functional brain studies in humans further support a role for insular cortex in complex auditory functions, with strong evidence for sensory responses in the posterior insula. Auditory responses in the posterior insula resemble those observed in Heschl’s gyrus^7^. The posterior insula is bilaterally activated when listening to speech, and this activation is believed to be sensory, rather than motor^8^. The posterior insula responds to speech coincidentally with the superior temporal gyrus, but in contradistinction is more active for self-generated speech than external speech^7^. The anterior insula, in contrast to the posterior insula, responds to the emotional contents of auditory stimuli, similarly to the amygdala^7,9^.

Animal studies also support a sensory role for the posterior insula. In awake squirrel monkeys, auditory neurons in the insula responded with latencies consistent with direct connections from the medial geniculate body (MGB). They were found in higher numbers in the posterior than in the anterior insula^10^. Injecting primate MGB with an anterograde tracer resulted in intense labeling of the junction between the temporal lobe and the insula (the parainsular field) throughout its extent^11^. In another vocal primate, rhesus monkeys, the posterior insula has been identified as a predominantly auditory region with neuronal representations of conspecific communication sounds that differed somewhat from the responses of neurons in auditory cortex to the same stimuli^12^.

These results in humans and in primates consistently share two striking properties. Auditory responses in the posterior insula have been shown to have short latencies and to have significant similarities with the auditory cortex. Yet, the posterior insula may also differ from auditory cortex, seemingly being responsible for complex aspects of vocal communication.

Much less is known about auditory processing in human and primate parietal somatosensory cortex. The human parietal operculo–insular region, where secondary somatosensory cortex and the posterior insula share a boundary, may participate in auditory–somatosensory cross-talk. The auditory function of this region is established, for example by the finding that its stimulation produces auditory hallucinations^13,14^. It has been proposed that the disruption of auditory–somatosensory integration in this region may result in phantom perceptions, such as tinnitus^15^. In some patients with somatosensory thalamic infractions, acquired auditory-tactile synesthesia may occur: patients reported that certain sounds produced an intense somatosensory tingling sensation. This sensation was accompanied by a threefold increase in the responses to sounds in the parietal operculum, the location of secondary somatosensory cortex^16^. Recent data further confirmed auditory responses in human somatosensory cortex^17,18^. In primates as well, a small portion of single neurons recorded in secondary somatosensory cortex respond to sound^19,20^.

There is evidence for functional cross-talk between the insular and parietal cortex in taste perception, nociception, temperature, and somatosensation^21–26^. Indeed, rodent insular and parietal cortices are intracortically connected^22,23,27^.

This short survey shows that the posterior insula and the nearby somatosensory field provide an intriguing window into the role of secondary auditory fields. In rats and mice, the posterior insula is innervated by the MGB and contains auditory neurons ^25,28–33^. Also in rats, short-latency auditory responses were detected in both primary and secondary somatosensory fields^24,34–36^. In contrast, in mice the evidence for purely auditory activity in secondary somatosensory cortex is weak^37–40^.

Importantly, in rats, the auditory-responsive area in the posterior insula is anatomically separated from other auditory fields, unlike in mice^25,28,30^. The clear presence of auditory responses in rat insular and somatosensory cortices, together with their anatomical separation from the rest of rat auditory cortex, make the rat into an excellent model for studying these fields.

In the current work, we studied the anatomical and functional organization of the auditory activity in the posterior insula and secondary somatosensory cortex in anesthetized rats. We recorded neuronal responses using standard electrode arrays as in previous studies^12,28,30,20^, as well as using state-of-the-art Neuropixels probes. High-density silicon probes allowed us to delineate an auditory field that spans the border between the posterior insula and secondary somatosensory cortex. We characterized the auditory response properties to broadband noise and pure tones. As a test case of more complex processing, we also studied deviance detection in these fields.

## Results

Extracellular spiking activity was recorded in fifteen rats from the left posterior insular cortex (Ins) and the area of the secondary somatosensory cortex (S2) adjacent to Ins dorsally. In addition, auditory units were also recorded in the primary auditory cortex (A1), secondary auditory fields (AC), medial geniculate body (MGB), and inferior colliculus (IC), in a separate group of five rats for each region, using the same set of sounds. In the rat brain atlas, A1 corresponds to Au1, and AC corresponds to both AuD and AuV^41^.

### Location of auditory units in insulo-parietal (Ins/S2) auditory field in the rat

Recordings of neuronal activity were obtained using two types of electrodes: monopolar tungsten electrodes (n = 7) and Neuropixels silicon probes (n = 8, 23 independent brain penetrations) (Fig. 1). Neuropixels probes facilitated high-yield recordings of neuronal activity from multiple units in Ins and S2 with each penetration.

**Figure 1.**
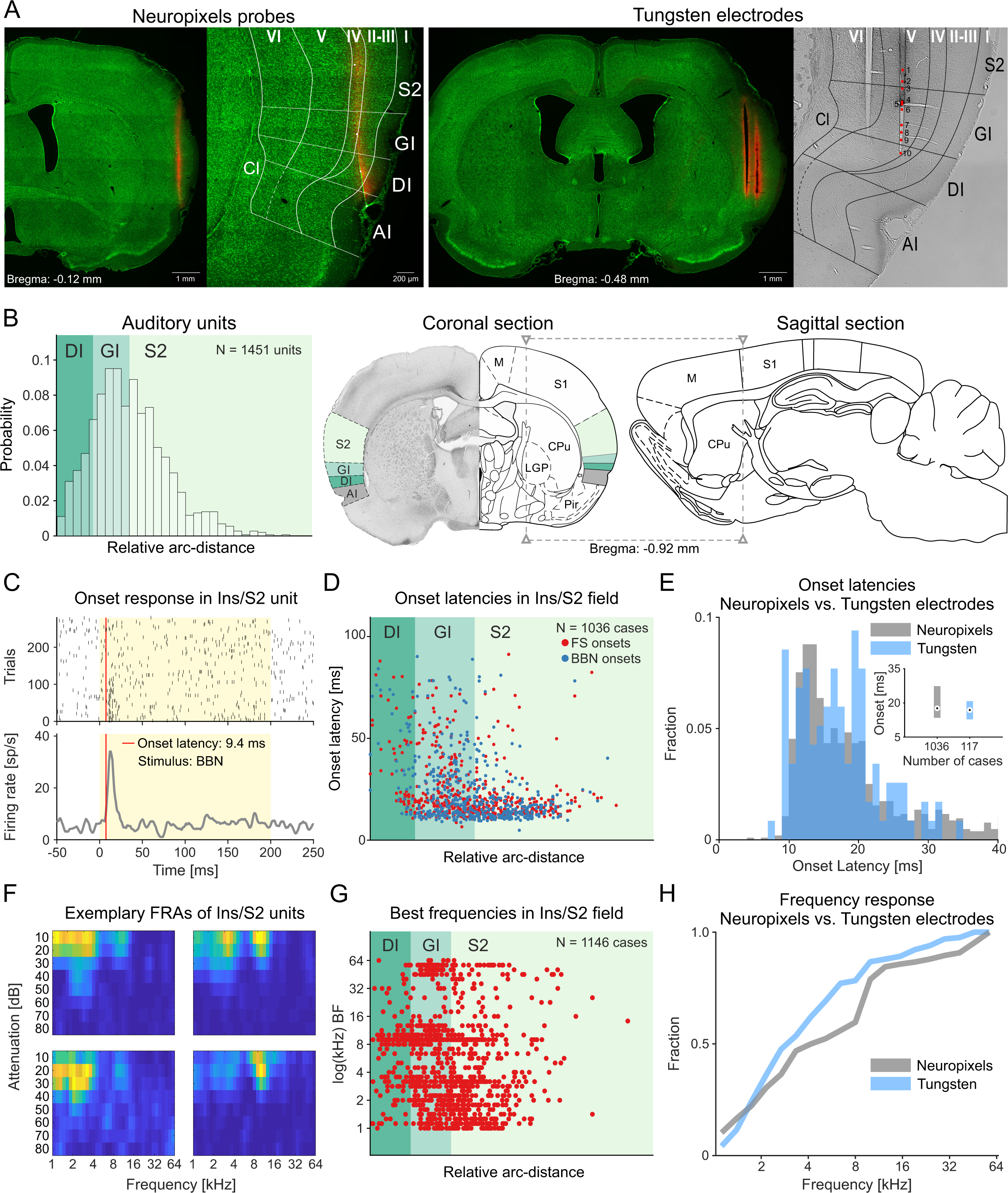
Location, onset latencies, and frequency tuning of auditory neurons in Ins/S2 cortical field. **(A)** Representative coronal brain sections with electrode tracks passing through Ins and S2 for Neuropixels (left side) and tungsten electrodes (right side). Images include an enlarged view of the region of interest (ROI) with division into Ins subregions (GI, DI, AI) and cortical layers. The recording locations for tungsten electrodes were marked with red dots on the bright field image of ROI. Brain sections were stained with fluorescent green Nissl’s stain, and electrodes were coated with DiI for better identification of the track. **(B)** Likelihood of occurrence of auditory neurons recorded with Neuropixels probes across Ins subregions and S2. The majority of auditory units were found in the GI and ventral portions of S2. On the right, we depicted approximated lateral and antero-posterior location of the Ins/S2 auditory field in rat brain. **(C)** A raster plot and histogram showing the representative unit response to broadband noise, characterized by a typical short onset latency (9.4 ms). **(D)** Onset latencies for all units recorded with Neuropixels probes in response to frequency sweeps (FS) and broad band noise (BBN). Consistently, large portion of GI and S2 units had short onset latencies <20 ms. **(E)** Comparison between normalized onset latencies of units recorded with Neuropixels vs. tungsten electrodes showed similar results for both techniques. **(F)** Frequency response arrays (FRAs) of exemplary units recorded in Ins/S2 field showing preference towards frequencies <16 kHz. **(G)** Best frequencies (BFs) for all units recorded with Neuropixels probes. The BFs <16 kHz were dominating. **(H)** Comparison between normalized BFs of units recorded with Neuropixels vs. tungsten electrodes showed similar results for both techniques. Abbreviations: I – cortical layer 1; II-III – cortical layers 2/3; IV – cortical layer 4; V – cortical layer 5; VI – cortical layer 6; AI – agranular insular cortex; Cl – claustrum; CPu – caudate-putamen; DI – dysgranular insular cortex; GI – granular insular cortex; LGP – lateral globus pallidus; M – motor cortex; Pir – piriform cortex; S1 – primary somatosensory cortex; S2 – secondary somatosensory cortex.

To determine the location of auditory units, we used the consecutive coronal brain sections covering the area with visible electrode tracks. Based on the histological material, we identified which neurons were recorded in the Ins and S2, their cortical layers, and the subregions of Ins: granular insular cortex (GI), dysgranular insular cortex (DI), and agranular insular cortex (AI) (Fig. 1A). The numbers of auditory and non-auditory units recorded in all rats across these structures are listed in Table 1 and supplementary Table S1. We drew the borders between Ins subregions based on the presence and morphology of granular layers and their relative position with respect to other brain structures (i.e., claustrum, endopiriform nucleus, piriform cortex, external capsule, anterior commissure, septum, ventricles, thalamic nuclei, and other structures depending on anteroposterior coordinates of given brain section). All histological reconstructions and divisions were made in accordance with the rat brain atlas^41,42^.

**Table 1.**
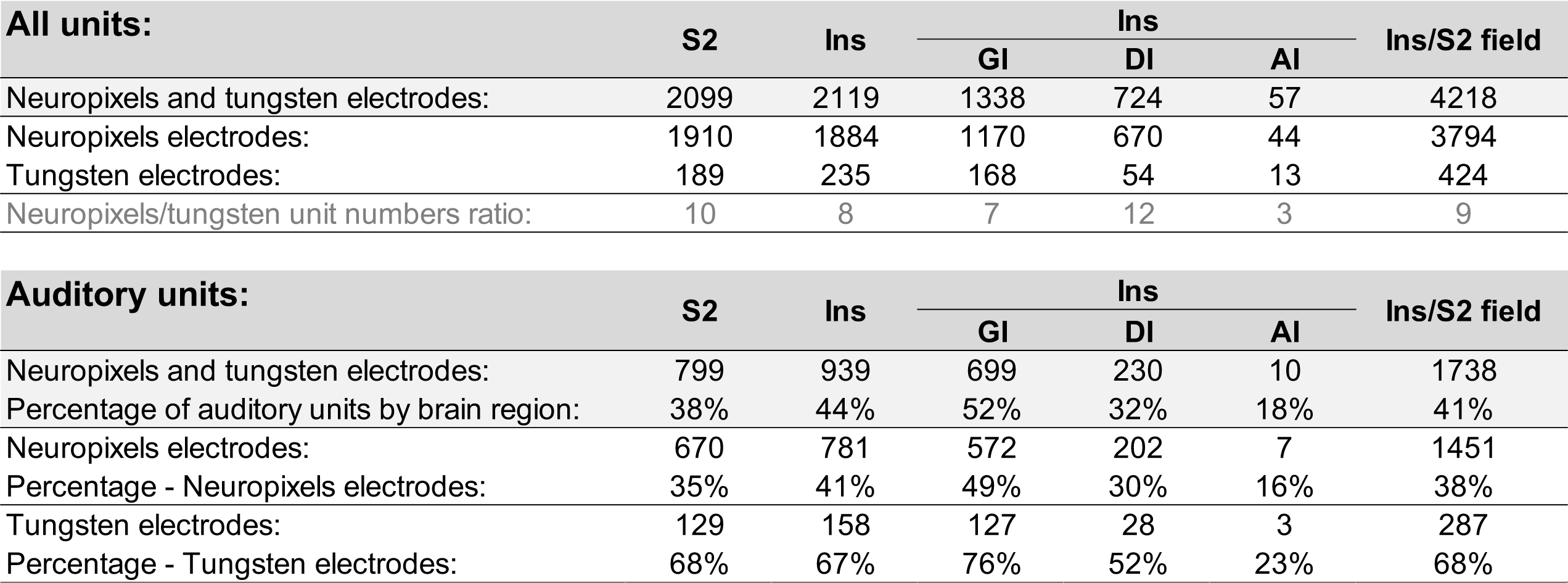
The numbers of all detected units and auditory cells in the secondary somatosensory cortex (S2) and insular cortex (Ins) were obtained using both Neuropixels and Tungsten electrodes. Counts are divided by Ins subregions: granular insular cortex (GI), dysgranular insular cortex (DI), and agranular insular cortex (AI), as well as for the whole parieto-insular field (Ins/S2). Additionally, the percentages of auditory units for Neuropixels and tungsten electrodes are reported. The lower percentage of auditory units in the neuropixels recordings are likely due to the much higher yield of these recordings, that include often units with very low firing rates, too low to demonstrate any statistical relationships with the sound presentations.

We recorded a total of 4218 units in the Ins/S2 field, out of which 1738 (41%) were classified as auditory neurons (see Methods for criteria). In the posterior Ins there were 939/2119 (44%) auditory units. The distribution of auditory units across Ins subregions was as follows: GI 699/1338 (52%); DI 230/724 (32%); AI 10/57 (18%). In S2, we recorded 799/2099 (38%) auditory units (Fig. 1B). The median anteroposterior coordinate of the electrode tracks was −0.48 mm posterior to bregma (max: +0.24 mm; min: −1.56 mm, n = 48 electrode tracks).

We compared these results with neural responses recorded previously in three main stations of ascending auditory system: the inferior colliculus (IC, midbrain), the medial geniculate body (MGB, diencephalon), and the primary and secondary auditory cortex (A1 and AC, telencephalon), each in five rats (Harpaz et al. 2021). The units were analyzed in the same way as those in Ins/S2. We recorded 2074/3983 (52%) auditory units in IC; 2298/5145 (45%) auditory units in MGB; and 2776/5619 (49%) auditory units in A1/AC (Table 2).

**Table 2.**
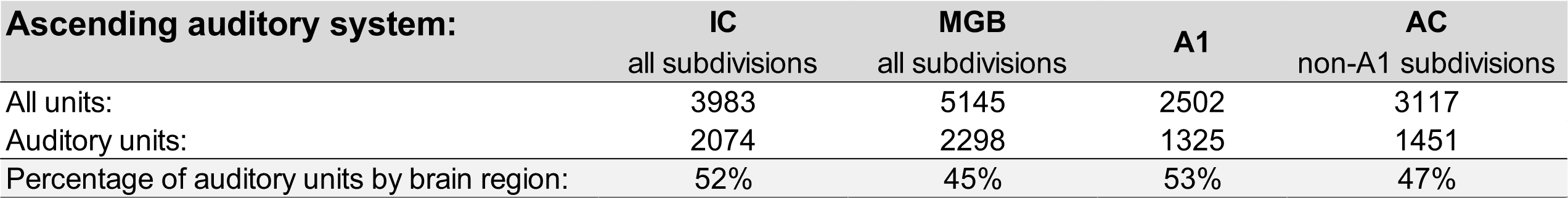
The numbers of all detected units and auditory cells in the structures of the ascending auditory system: the inferior colliculus (IC, all subdivisions), the medial geniculate body (MGB, all subdivisions) the primary auditory cortex (A1) and surrounding fields (AC, non-A1 subdivisions: Aud/Auv). The percentage of auditory cells per tested region is provided.

### Onset latencies

The majority of auditory units recorded in the Ins/S2 field had short onset latencies (median: 17.6 ms, IQR: 13.4 - 26.1 ms). This includes onset latencies to broadband noise (median: 16.8 ms, IQR: 12.3 - 27.3 ms) and to pure tones (median: 18.4 ms, IQR: 14.0 - 25.1 ms) (Fig. 1C and D). We observed similar distribution of onset latencies for units recorded in Ins/S2 field with tungsten and Neuropixels electrodes across all rats (Fig. 1E).

The onset latencies differed among the different anatomical subdivisions. Shorter latencies were observed in S2 and GI, while longer latencies were observed more ventrally in DI. In S2, the median latency was 15.2 ms (IQR 12.5 - 19.8 ms). In GI, the median latency was 19.4 ms (IQR 13.8 - 31.2 ms) and in DI, the median latency was 34.1 ms (IQR 20.4 - 56.5 ms). All differences were significant: both Ins regions had significantly longer median onset latencies than S2 (GI vs. S2: z= 7.9285, rank sum = 315898, p = 2.2e-15 and DI vs. S2: z = 10.5738, rank sum = 46629, p = 3.9e-26, two-sided Wilcoxon rank sum test), while units in the DI region had significantly longer latencies than units in the GI region of Ins (z= 6.9075, rank sum = 39483, p = 4.9e-12, two-sided Wilcoxon rank sum test). Minimum latencies in S2 and GI were similar, but GI had in addition a population of longer-latency neurons. DI lacked neurons with relatively short onset latencies (<20 ms) compared to GI and S2 (*χ*^2^= 70.9421, df = 1, p = 3.7e-17, *χ*^2^test).

We compared these latencies with those measured in IC, MGB and AC using the same methods (Fig. 2A and B). As expected, IC units typically had shortest onset latencies (median:11.2 ms, IQR: 9.6 - 13.1 ms) than MGB units (median: 12.8 ms, IQR: 11.0 - 15.5 ms). Surprisingly, auditory neurons in Ins/S2 field had significantly shorter onset latencies (median: 17.6 ms, IQR: 13.4 - 26.1 ms) than in A1 (median: 20.3 ms, IQR: 17.6 - 24.0 ms) (z = −9.4600, rank sum = 1229470, p = 3.1e-21, two-sided Wilcoxon rank sum test) and surrounding fields (median: 24.1 ms, IQR: 19.4 - 42.1 ms) (z = - 17.6550, rank sum = 987401, p = 9.3e-70, two-sided Wilcoxon rank sum test). Examples of representative neurons recorded in each of these regions are presented in Figure 3.

**Figure 2.**
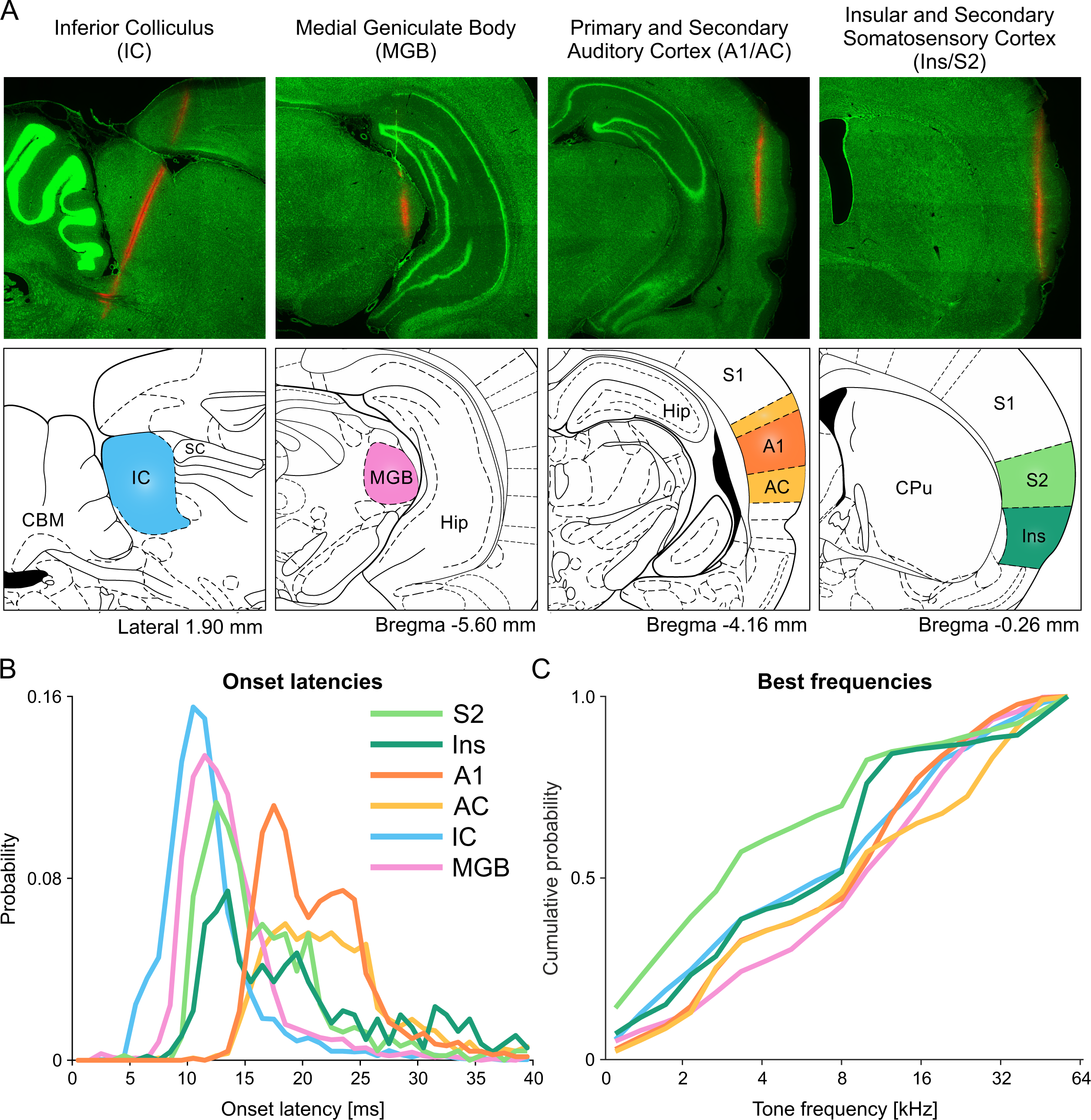
Onset latencies and BFs across structures of ascending auditory system and Ins/S2 field. **(A)** From left to right, we show representative fragments of sagittal and coronal brain sections with Neuropixels probes tracks in the inferior colliculus (IC), medial geniculate body (MGB), primary and secondary auditory cortices (A1/AC), and Ins/S2 field. Below each histology slide are schematic drawings depicting the borders of structures for chosen antero-posterior coordinates. **(B)** Likelihood of onset latencies for all units in tested brain structures depicting very short onset latencies of neurons in Ins/S2 field in comparison to A1 and AC auditory cortices. **(C)** Probability of BFs for all units in tested brain structures showing lower counts of units with BFs 16-32 kHz in Ins/S2 field compared to structures of ascending auditory system. Abbreviations: A1 – primary auditory cortex; AC – secondary auditory cortices; CBM – cerebellum; CPu – caudate-putamen; Hip – hippocampus; IC – inferior colliculus; Ins – insular cortex; MGB – medial geniculate body; S1 – primary somatosensory cortex; S2 – secondary somatosensory cortex; SC – superior colliculus.

**Figure 3.**
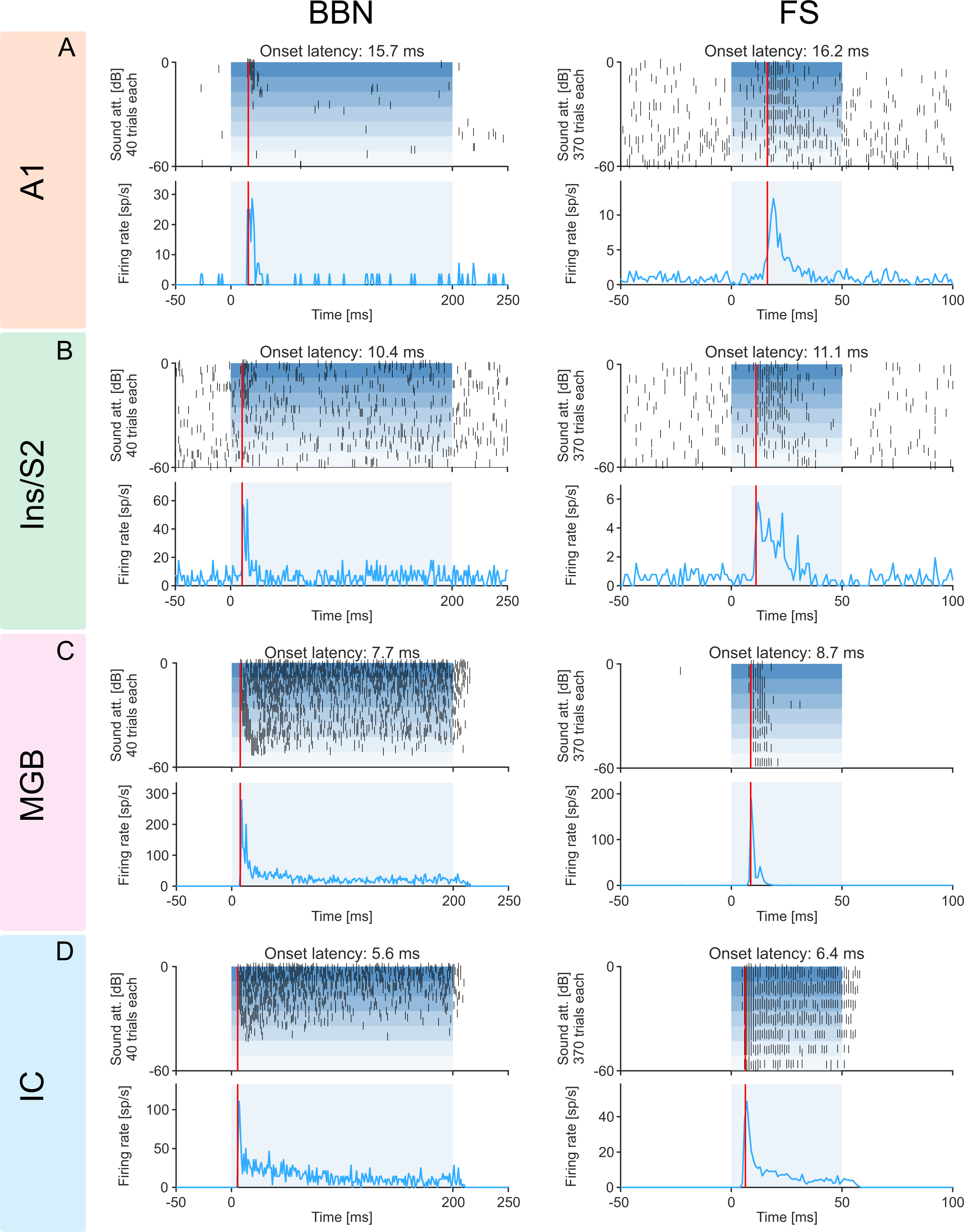
Exemplary responses of units to Broad Band Noise (BBN) and Frequency Sweeps (FS). Representative neuronal responses to BBN (280 trials, 10 for each sound level, left column) and FS (370 trials, 10 sound levels for 37 frequencies, right column) of the units recorded in **(A)** the primary auditory cortex (A1), **(B)** the insulo-parietal field (Ins/S2), **(C)** the medial geniculate body (MGB), and **(D)** the inferior colliculus (IC). In each panel, raster plots are presented with corresponding mean firing rates below. The red vertical line represents onset latency and shading stimulus duration. The examples were chosen to illustrate the general trend of average latencies getting longer in the following order: IC → MGB → Ins/S2 → A1.

### Frequency tuning

Frequency tuning was evaluated based on the responses to pure tones (1–64 kHz, 6 tones/octave; Fig. 1F-H and 2C). Neurons recorded in DI, GI and S2 were preferably tuned to tones of frequencies below 16 kHz when compared to A1 and surrounding fields (*χ*^2^ = 136, *df* = 1, p=2.0e-31), as depicted in representative example (Fig. 1F) and for all units (Fig. 1G and H). Median best frequency was 4.8 kHz, IQR: 2.0 - 10.1 kHz (in A1 and surrounding fields: median 9.0 kHz, IQR 20.2 – 28.3 kHz).

### Stimulus-Specific Adaptation (SSA)

SSA is the decrease in the responses to a commonly-presented stimulus (‘the standard’) which is not, or only partially, generalized to another, rare frequency (‘the deviant’). Neurons throughout the auditory system show stimulus-specific adaptation (SSA). To study SSA, two tone frequencies were selected within the frequency range of the units recorded in each site, and each of these frequencies was tested in six different conditions (see Methods for details). Three of these conditions consisted of sequences composed of the two frequencies only: standard (95% of tone presentations in the sequence), deviant (5% tone presentations), and equal (50% tone presentations). The ‘rare’ (with inter-tone intervals of many seconds) provided estimates of the least adapted responses. The other two conditions were multi-tone sequences in which each of the two tested frequencies were presented 5% of the time. In one of these sequences the other tones covered a narrow frequency range (‘diverse-narrow’) and in the other sequence they covered a wide frequency range (‘diverse-broad’). Unlike in oddball sequences, where deviant sounds violate the expectation for the next, more frequent standard sound to appear, in the multi-tone sequences no expectations can be formed for the next tone frequency. The two multi-tone sequences differ however in the amount of across-frequency adaptation that may occur^43^.

To quantify how presentation probability affects tone response (Fig 4A and B), we calculated the common contrasts (CSI) between the deviant and standard responses to characterize the average effect of adaptation for this specific pair of frequencies:

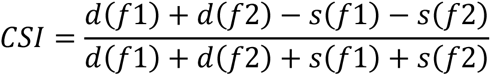

**Figure 4.**
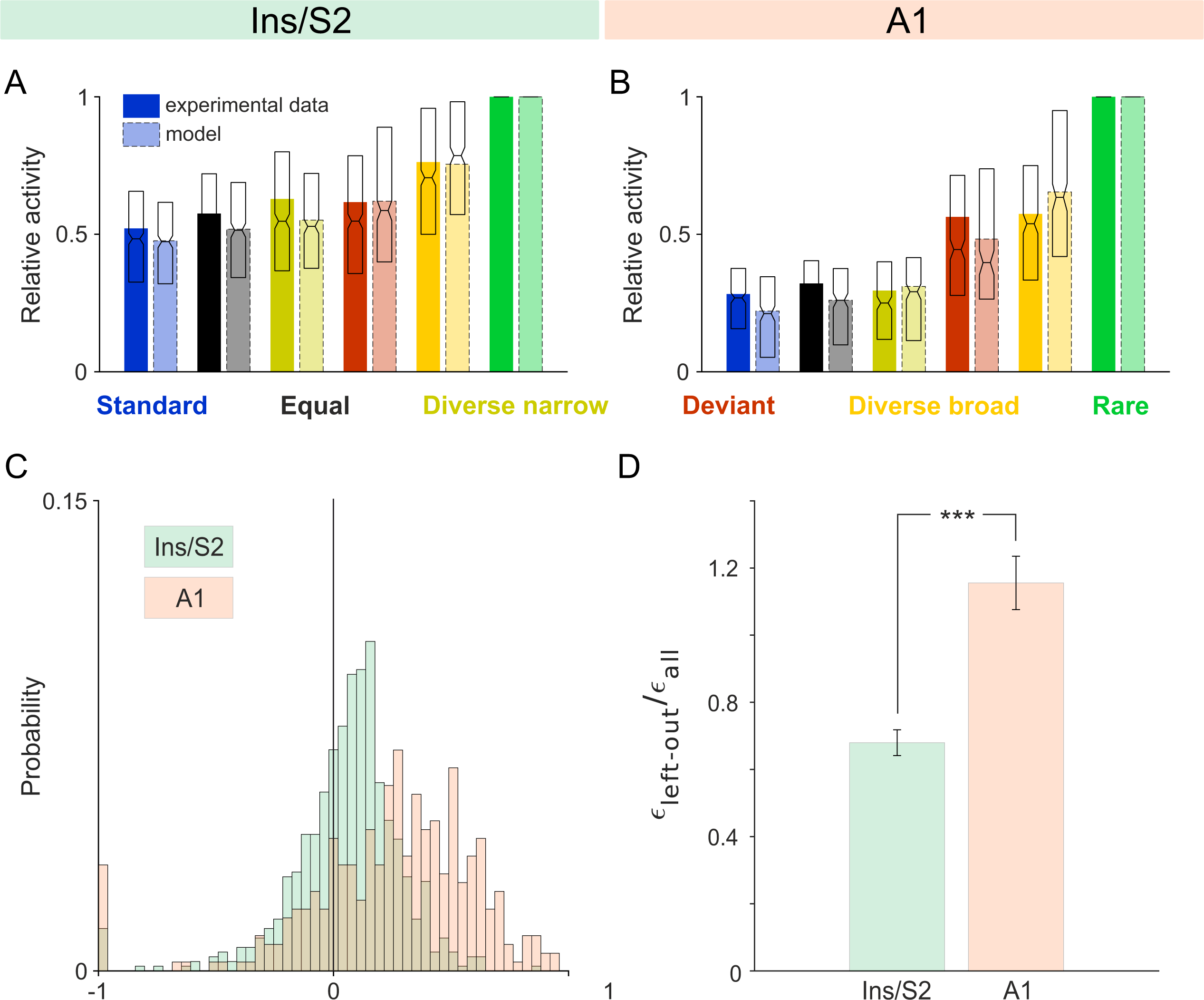
Stimulus-specific adaptation: Ins/S2 auditory field vs. A1. Panels **(A)** and **(B)** show the mean responses (bars) and median + IQR values (overlayed boxplots) of MUA (solid-color bars) and the model-predicted responses (faded-color bars outlined by dashed lines) for Ins/S2 and A1, respectively. As previously reported, in A1, responses to deviant tones in diverse broad and oddball sequences were indicative of deviance sensitivity due to larger-than-expected deviant responses. In contrast, in Ins/S2, diverse broad responses exceeded deviant responses, demonstrating better alignment with the model. In **(C)**, overlaid histograms of CSI for Ins/S2 and A1 are presented. Overall, we observed a higher proportion of positive CSI values for units in A1 than in Ins/S2, suggesting a frequency-independent response component to a stimulus when deviant. **(D)** Deviant prediction error for Ins vs. A1. Plotted here is the mean ± SEM of the normalized error values obtained from the leave-one-out analysis of the NFAC model. Error values are calculated by the difference of the normalized predicted deviant response from the model fitted on all conditions excluding the deviant response, and the measured normalized response. Afterwards we normalize the error value for each unit by dividing it by the average error in the model predictions for all other responses, when fitted on the complete set of responses. A1 normalized error values for deviant response when left out are significantly higher than Ins/S2 (***p < 0.001, Wilcoxon rank sum test). Such that deviant responses in A1 are poorly accounted for by simple adaptation profiles, implying deviance sensitivity, when compared to Ins/S2 responses.

where *d(fi)* and *s(fi)* represent the average responses to frequency fi when it was deviant and standard, respectively. The average CSI in Ins/S2 (mean: 0.054, ± 0.009 SE) was significantly smaller than in A1 (mean: 0.242, ± 0.019 SE, t=-9.8335, df = 1019, p=7.3e-22, two-sample t-test; Fig. 4C).

We also calculated the frequency-specific index SSA index (SI),

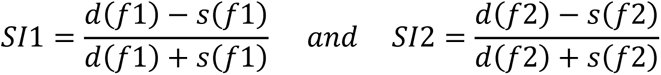

Overall, among Ins/S2 neurons there was a smaller number of cases where both contrasts were positive (317/666, 48%) than in A1, where this effect was stronger (194/355, 55%; *χ*^2^ = 4.99, *df* = 1, p = 0.025). Thus, Ins/S2 units showed a weaker effect of stimulus probability on their responses than A1 units.

To quantify deviance sensitivity and explain the adaptation profiles in Ins/S2 and A1, we used a previously established model based on adaptation in narrow frequency channels (ANFC)^43^. The model assumes that the neural responses to any frequency depend on the available resources in a narrowly tuned frequency channel centered on it. These resources are depleted by tone presentations according to how far they are from the center of the frequency channel, and recover exponentially between stimulus presentations. The model has 2 parameters, one for the width of the frequency channels, and the other for the recovery time constant of the resources between tone presentations (see methods). The width of the frequency channels determines the effects of presentations of a tone of one frequency on the responses to tones of another frequency through the ‘adaptation load’ – the average amount of resources depleted within the channel centered on the test frequency, that differ for the different sequences used here^43^ (see Methods for detail). The model was fitted to units in Ins/S2 and A1, and predictions fitted well to the measured responses (Fig. 4A and B).

We used the model to analyze the deviance sensitivity of neurons in Ins/S2 relative to neurons in A1 using the method of Taaseh et al.^43^. Deviance sensitivity is tested by comparing the responses to the same frequency in the diverse broad and in the deviant conditions. In both conditions, the test frequencies are played with the same probability (5%). However, in the diverse broad condition, all frequencies are presented with low probabilities, so that none is special (no ‘deviance’). This is in contrast with the deviant condition, in which the standard may generate expectations that are violated by the presentations of the deviant tone^44^. Thus, deviance sensitivity expresses itself as responses that are larger to deviants than to the same sounds in the DB sequence.

On the other hand, the ANFC model predicts responses that are smaller to deviants than to the same sounds in the DB sequence, because for the deviant, all other tone presentations are the standard tone which is about half an octave away, while in the DB sequence many of the other tone presentations are farther away from the tone of interest (see ^43^ for a quantitative treatment). In A1, DB responses and deviant responses were similar (t = −0.33, df =354, p = 0.74), reproducing previous results^43,45,46^ (Fig. 4B). On the other hand, in Ins/S2, DB responses were significantly larger than deviant responses (t = −9.03, df = 665, p = 1.7e-18). Furthermore, in Ins/S2, the normalized differences between the responses to DB and deviant were significantly larger than the differences observed in A1 (t = 2.8, df = 1014, p = 0.0057), better fitting the model (Fig. 4A).

We have previously interpreted the similarity in responses to deviant and DB in A1 as indicating the presence of deviance sensitivity, since the responses to the deviant condition were larger than expected by the ANFC model, suggesting the presence of an additional response component that was sensitive to deviance. To further test deviance sensitivity in our data, we ran a leave-one-out analysis, in which we fitted the model using the responses to all tone sequences except for the deviant condition, and compared the predicted responses in the deviant condition to the measured ones. There were significantly larger errors in predicting the deviant response of units in A1 than for units in Ins/S2 (Fig. 4D, z = −6.8436, rank sum = 142545, p = 7.72e-12, two-sided Wilcoxon rank sum test), supporting the conclusion that units in A1 show more deviance sensitivity than neurons in Ins/S2.

## Discussion

We describe here basic firing properties of auditory neurons in the insular and secondary somatosensory cortices in anesthetized rats. Auditory neurons were present in the posterior insular cortex, particularly in the granular insular cortex, extending medially to the adjacent ventral part of the secondary somatosensory cortex. In the rest of the discussion, this contiguous area will be referred to as the insulo-parietal field.

Units throughout the insulo-parietal field shared some response properties: (1) We describe a medio-lateral gradient of response latencies. In the medial part of the field, units had on average shorter onset latencies to broad band noise and pure tones than more laterally, with the longest latencies in that part of the auditory field that lies in dysgranular insular cortex (DI). Remarkably, some neurons had shorter onset latencies than the shortest latencies recorded using the same methods in the primary and secondary auditory cortices. (2) The insulo-parietal auditory units were found to be preferentially tuned to relatively low frequencies (<16 kHz). (3) The habituation profiles of neurons in the insulo-parietal field were subtly different from those found in the primary auditory cortex, with deeper adaptation and less deviance sensitivity.

These observations establish the presence of an ultra-fast auditory information stream through the insulo-parietal field, in parallel to the usual primary pathway to primary auditory cortex and its surrounding fields.

### Insulo-parietal auditory field in rodents

Most studies of the parcellation of auditory fields in rats ignored the more anterior areas studied here^47,48^. Nevertheless, previous research has identified neurons in the insular and somatosensory cortices that are responsive to sound. In rats, auditory activity in the insular cortex has been reported in several studies. For example, Rodgers et al. (2008)^25^ used epipial recordings to identify the cortical area that responds to sound. In rats, this area is located in the posterior insula and is anatomically separated from the auditory cortices in the temporal lobe by the insular somatosensory field^34,25^. In another study, Kimura et al. (2010)^28^ used precise but low-yield recording and tracing methods, followed by histological analysis, to establish the presence of auditory responses in the posterior insular cortex. The anterior-posterior location and extent of this field is largely consistent across these two studies and ours (Rodgers et al. (2008)^25^: +0.5 to −1.5 mm posterior to bregma; Kimura et al. (2010)^28^: +0.3 to −1.3 mm posterior to bregma; Current study: +0.24 and −1.56 mm posterior to bregma).

The current report extends these two studies, and establishes the presence of auditory neurons not only in the insular cortex but also more medially, in the ventral portion of the secondary somatosensory cortex. Earlier studies have reported the presence of multimodal somatosensory-auditory units on the border between the parietal and temporal lobes^34,24,36^. Very few studies have described neuronal responses to sounds in somatosensory cortices of rats^24,35^, and of mice^37–40^. Rodgers et al. (2008)^25^ attributed auditory function exclusively to the insular cortex, but their methods could not accurately distinguish borders between morphologically similar cortices. In Kimura et al. (2010)^28^, some auditory units were located on the putative border between these two regions, yet were attributed by the authors to the insular cortex (e.g. Fig. 3 in Kimura et al. (2010)^28^. The small sample size in that study may have contributed to its focus solely on the insular cortex.

The current study used both tungsten electrodes and neuropixels probes to record an extensive neuronal population, demonstrating the presence of auditory units in both insular and secondary somatosensory cortices. Neuropixels probes provided several advantages for functional anatomy studies. They allowed us to record simultaneously across both areas, record significantly more units, and minimize experimenter bias during the search and selection of units for recording. Thorough histological analysis ensured the accuracy and reliability of our results, establishing the presence of auditory neurons in the ventral part of secondary somatosensory cortex in addition to the previously reported auditory neurons in insular cortex.

Among rodents with lissencephalic brains, the separation of the insulo-parietal field from the more posterior core auditory fields appears to be well-documented only in rats^34,25,49^. In mice, an insular auditory field exists as well, but is continuous with the anterior auditory field (AAF) in the ventrorostral direction^29–33^. Thus, Sawatari et al. ^29^ identified and characterized the insular auditory field in mice using wide-field calcium imaging followed by histological analysis. Guo et al. (2012)^30^, using electrophysiological mapping in anesthetized and awake animals, made similar observations, confirming that the insular auditory field in mice is continuous with the AAF.

The anatomical separation in rats between the insular auditory field and other auditory fields provides a distinct advantage relative to mice, where the insular auditory field is continuous to other auditory fields, making it harder to distinguish between them. Rats therefore offer a valuable model for investigating the insular auditory field.

### Auditory inputs to insular and secondary somatosensory cortices

Evidence for thalamic inputs from the ventral, primary, division of the MGB to insular cortex has been reported in cats^50^. In mice, the ventral division of the MGB provides direct input to the insular cortex. Notably, those thalamic neurons projecting to the insular cortex constitute a distinct population from those projecting to either the primary auditory cortex or AAF^32^.

In rats, the posterior insular cortex is innervated by non-lemniscal auditory thalamic nuclei, including the medial division of the MGB and the suprageniculate thalamic nucleus, as well as the non-specific posterior intralaminar thalamic nucleus^51–53^. Evidence for direct inputs from the ventral division of the MGB to the insular cortex in rats is lacking. In addition to direct auditory thalamic inputs, the posterior insular cortex receives inputs from the ventral auditory field (VA) which is largely innervated by the dorsal division of the MGB^54^.

The ventral area of secondary somatosensory cortex, where we found auditory neurons in rats, is less explored in terms of auditory inputs, particularly those originating in the thalamus. We review here the known facts, and subsequently delineate its inputs to shed light on its functional relevance. The ventral part of the secondary somatosensory cortex has been suggested to be a separate field, containing a complete representation of the contralateral body surface. This has been first demonstrated using microelectrode unit recordings in gray squirrel^55^, where two complete representations of the contralateral body surface have been delineated in secondary somatosensory cortex: the second somatosensory cortex proper (SII) and a parietal ventral area (PV). Subsequent studies in rats followed the division established by Krubitzer et al. (1986)^55^, similarly identifying an SII and a PV subregions ^56–58^. Remple et al. (2003)^57^ described SII as having an upright body representation located just lateral (ventral) to the primary somatosensory cortex, while PV has an inverted representation and is situated laterally (ventrally) to SII.

SII and PV receive auditory inputs in a number of species. In cats, an area of secondary somatosensory cortex analogous to PV receives projections from the medial division and the suprageniculate nuclei of the MGB^59^. Interestingly, in mice, a partial overlap between the parietal ventral area (PV) and secondary auditory fields, including the insular auditory field, have been described^60^. Takemoto et al. (2014)^32^ previously demonstrated that the mouse insular auditory field, which overlaps with the PV, is directly innervated by the ventral division of the MGB. Thus, while evidence for innervation of PV by the ventral division of the MGB is lacking in rats, it appears to exist in mice.

In our study, we recorded neural activity in the most ventral part of secondary somatosensory cortex neighboring granular insular cortex, as defined by Paxinos and Watson^41^. This region is likely to correspond to the head representation within the PV region of Remple et al. (2003)^57^.

Summarizing, the rat insulo-parietal field likely receives auditory inputs from the non-primary MGB, including the medial division of the MGB and the suprageniculate nucleus. Definite information about the existence (or lack thereof) of input from the ventral division of the MGB is lacking.

### Short onset latencies

One of the most surprising results of this paper is the observation of units in the insulo-parietal field whose onset latencies to broad band noise and pure tones were shorter than the shortest response latencies recorded by us in the primary and secondary auditory cortices, using exactly the same methods.

Consistent with our data, Rodgers et al. (2008)^25^ used epipial electrode recordings in rats and reported that poststimulus latencies of the P1 and N1 peaks in the insular auditory field were significantly shorter than latencies for the same components in the AAF, suggesting that the insular auditory field does not rely on intracortical projections from the AAF for access to auditory input. Other studies using similar methodologies have also reported low latencies. Kimura et al. (2010)^28^ reported the lowest onset latency of about 10 ms (on average 12.7-19.8 ms) in rats, comparable to the shortest onset latencies in A1. Our data, however, are the first demonstration of spiking response latencies in the insulo-parietal field that are shorter than those found in A1.

In mice, as well, responses in insular cortex are remarkably fast. For example, Guo et al. (2012)^30^ observed latencies of approximately 10 ms on average in the insular auditory field of mice. Similarly, Sawatari et al. (2011)^29^ observed the fastest cortical calcium responses to 4 kHz tone in the insular auditory field. Subsequent studies by the same group demonstrated that this cortical region is directly innervated by the ventral division of the MGB^32,60^. At least in mice, the short response latencies in the insulo-parietal field can be accounted for by these connections.

In the rat, evidence for innervation by the ventral division of the MGB is lacking. The posterior insular cortex is directly innervated by the medial division of the MGB, suprageniculate thalamic nucleus, and non-specific posterior intralaminar thalamic nucleus^51–53^. Of these, the medial division of the MGB has the largest neurons across the auditory thalamus^61^. These neurons receive inputs from the lateral and rostral cortex of the inferior colliculus^62^ but are also directly innervated by axons from the dorsal cochlear nucleus^63^. It is possible that the pathway from DCN to the medial MGB, bypassing the IC, may be the source for the short latency responses in the insulo-parietal field^64^.

### Tuning to a low frequency range

In both mice and rats, the insular field has been reported to over-represent low frequencies. In mice, insular auditory field was most responsive to 4 kHz tones compared to 8, 16, and 32 kHz^29^. It is possible that the insulo-parietal field is involved in processing hetero- and conspecific vocalizations, including distress calls. For example, when mice are stressed they often emit a vocalization with a fundamental frequency of about 4 kHz. When sonic calls were played back to rats and mice, it was found that these vocalizations elicited approach behavior in both species, although only in the presence of a live mouse^65^. Rats also communicate in the audible range (<16 kHz), and these low-frequency calls are often associated with pain-related behavior or produced in response to noxious stimuli^66^. Furthermore, vocalizations of species that prey on small rodents typically involve energy below 4 kHz^67,68^.

### Response habituation in insulo-parietal field

We observed a relatively simple adaptation profile in auditory neurons located in the insulo-parietal field. As in A1, the responses to test tones increased in strength in the following order across all tested conditions: standard → equal → diverse narrow → deviant → diverse broad → rare.

As in A1, tone frequencies close to the test tones seem to cause more adaptation than tones farther away^43,69–71^. However, responses were substantially reduced in all sequences relative to the least adapted rare condition, with smaller differences between the different conditions than found in A1. In particular, SSA was overall weaker in the insulo-parietal field than in A1, with deviant responses substantially more similar to standard responses than in A1.

The relative weakness of the deviant responses in the insulo-parietal field is particularly striking in comparison to the responses evoked by tones of the same frequency in the diverse broad condition. This is best illustrated by comparing the differences between responses to deviant and DB, which were significantly larger in Ins/S2 than in A1. Indeed, a simple adaptation model predicted better the responses to deviant tones in the insulo-parietal field than in A1. We have previously interpreted the discrepancy between the model and deviant responses in A1 as suggesting that the deviant responses showed true deviance sensitivity^43,45,46^. In the insulo-parietal field, in contrast, the responses to tone sequences seem to be largely governed by adaptation, without true deviance sensitivity. Nieto-Diego and Malmierca (2016)^48^ suggested that SSA to tones is stronger and develops more rapidly in non-primary auditory fields than in primary fields^72^. Our results here (as well as in ^46^) suggest the opposite: deviance sensitivity seems to be less pronounced in secondary auditory fields, largely because a stronger adaptation of the deviant tones by the accompanying standards. Further studies are required to resolve this issue.

### A fast pathway for the processing of communication sounds?

The short latencies of the insulo-parietal auditory neurons in rats indicate the presence of a fast stream of auditory information to cortex, parallel to the leminscal pathway through A1. Our results suggest the presence of multiple, parallel information pathways from the thalamus to the cortex. It is tempting to hypothesize that this pathway is specialized for processing communication sounds. These findings are particularly intriguing in view of the recent description of a parallel thalamo-cortical circuit for the processing of speech in humans^73^.

## Materials and methods

### Animals

Thirty female Sabra rats (240-325 g) that were naive to the experiments were used from Envigo LTD, Israel. The animals were housed in pairs per cage and had unrestricted access to water and standard rodent food prior to the experiments. The rats were maintained in a temperature and humidity-controlled room on a 12-hour light/dark cycle with lights on from 7:00AM to 7:00PM. The experiments were carried out in compliance with the regulations of the ethics committee at The Hebrew University of Jerusalem, an institution that is accredited by the Association for Assessment and Accreditation of Laboratory Animal Care (AAALAC).

### Electrodes and target brain structures

Extracellular recordings were conducted in the posterior insular cortex (Ins) and secondary somatosensory cortex (S2) using two types of electrodes: tungsten electrode arrays (in 7 rats) and Neuropixels silicon probes (in 8 rats). These data were compared to neuronal activity recorded in three structures of the ascending auditory system using Neuropixels electrodes in additional 5 rats per structure: primary and secondary auditory cortices (A1 and AC), medial geniculate body (MGB), and inferior colliculus (IC). The responses to other stimuli collected in these latter experiments have been previously published^74^ (tone clouds).

#### Tungsten electrodes

Two types of custom-built arrays were constructed using monopolar tungsten electrodes (0.6 MOhm, glass-coated, Alpha Omega Engineering, Nazareth, Israel). These arrays were built with 3 or 9 tungsten electrodes and were used in 2 and 5 rats respectively. The spacing between electrodes in the 3-electrode arrays was 300-500 µm and in the 3 × 3 electrode arrays it was 400-600 µm. The arrays were mounted on custom-made microdrives, which allowed for the gentle removal of the electrodes from fixed brain tissue after transcardial perfusion in order to preserve the shape of the electrode tracks in the brain tissue for precise histological reconstructions. Before each experiment, the electrodes were covered with multiple layers of orange-red fluorescent dye (DiI, ThermoFischer Scientific, Molecular Probes™, Eugene, OR, USA) dissolved in absolute ethanol (J.T.Baker, Deventer, Netherlands) and dried. Before and after the experiments, the electrodes were cleaned by washing them with trypsin solution 10X from porcine pancreas (Sigma-Aldrich, St. Louis, MO, USA) and purified water.

#### Neuropixels electrodes

Phase IIIA Neuropixels silicon probes (Imec, Leuven, Belgium) were used. A thin and flexible ground cable was soldered to the electrode’s PCB. The ground and reference contacts were shorted during the experiments. The ground cable was connected to a low-impedance reference contact on the head of the rat through a single gold-plated connector (MILL-MAX MFG. CORP., New York, USA). The electrodes were washed with trypsin solution 10X and rinsed with purified water before and after each experiment, and then placed for about 1 hour in a 4% solution of tergazyme in purified water (Alconox, Jersey City, NJ, USA). Finally, the electrodes were carefully washed in purified water. The underside of the silicon substrate of the electrode was coated with DiI dissolved in a 70% isopropanol solution in water (Sigma-Aldrich, St. Louis, MO, USA) and dried before use.

### Surgery

Rats were anesthetized with an initial intramuscular injection of ketamine hydrochloride (approximately 40 mg/kg, Clorketam 1 g/10ml, Vetoquinol, Lure, France) and medetomidine hydrochloride (approximately 0.1 mg/kg, Cepetor 1 mg/ml, Burgdorf, Germany). The surgical level of anesthesia was verified by the absence of pedal withdrawal and corneal reflexes, and by a slow, steady breathing rate. The animal’s head and neck were shaved, and body temperature was regulated using a closed-loop heating system controlled by a rectal probe (FHC, Bowdoin, ME, USA). The animal was placed on its back, and an incision was made on the neck above the trachea. The trachea was exposed by retracting fat tissue and muscle using blunt curved tweezers, and a transverse cut was made in the trachea. A stainless steel tube with blunted edges, covered at its end with a 4 mm long piece of polyethylene tubing (16G, made of injection needle 1.6 × 40 mm; Becton, Dickinson &Co. Ltd., Drogheda, Ireland) was inserted into the trachea. The metal tube was tied with a silk suture (Braided Silk Suture 6-S, Teleflex medical, Coventry, CT, USA) to the trachea above the thicker polyethylene part. Next, the animal was positioned on its ventral side, and the head was mounted in a custom head holder designed for auditory experiments, holding the animal by the maxilla. The holder was attached to the stereotaxic apparatus (David Kopf Instruments, CA, USA). The free end of the metal tube inserted into the trachea was attached to the tubing of the anesthetic machine with a time-cycled volume ventilator (Hallowell EMC, Pittsfield, Ma, USA). The animal was ventilated with 100% oxygen. The breathing parameters and CO2 level in exhaled gases were monitored using a capnograph (MicroCap, Oridion Medical 1987 Ltd., Brussels, Belgium). One hour after the injection of medetomidine and ketamine, 1% of halothane (Sigma-Aldrich, St. Louis, MO, USA) was added to the inhaled oxygen using a calibrated vaporizer (Matrx™ VIP 3000, Midmark Corporation, Versailles, OH, USA). After another half an hour, the halothane level was increased to 1.5% and adjusted individually if needed. The level of anesthesia was kept at a stable level throughout the entire experiment, and the CO2 level was maintained between 35 and 45 mmHg. The level of anesthesia used allowed for ventilation without muscle relaxation.

A longitudinal incision of the skin on the head (2-3 cm) was made, and the dorsal and left lateral surfaces of the skull were exposed. The connective tissue covering the bones was removed, and the bones were cleaned with sterile saline. Once the surface of the skull was clean and dry, a target reference point for the chosen recording site was marked on the bone (Ins/S2 - AP: −1.0 mm, ML: −6.1 mm from bregma; A1/AC - AP: - 5.1 mm, ML: guided by landmarks on temporal and parietal bones; MGB - AP: −5.6 mm, ML: −3.5 mm, IC - AP: −7.1 mm, ML: −2.0 mm). The electrode was inserted into the brain under an angle of 20 degrees to the vertical axis, one penetration was perpendicular to horizontal plane with the following coordinates: AP: −8.0 mm, ML: −2.0 mm. The soft tissue and bones, except around the implantation site, were covered with acrylic glue and acrylic dental cement (R21, AXIA, Seoul, Korea; Coral-fix, Tel Aviv, Israel). A well for saline was shaped around the implantation site using acrylic dental cement. A bare silver wire soldered to a thin, flexible ground cable was attached firmly with acrylic glue near the posterior edge of the implantation site (bare silver wire 0.01 inch, A-M Systems, Everett, WA, USA). The craniotomy was created by drilling (square approx. 3 × 3 mm).

### Tungsten electrode array insertion

The dura was gently resected. The electrodes were slowly (less than 10 µm/s) inserted vertically into the brain tissue until auditory units were detected (broad-band noise stimulus) using a single-axis motorized micromanipulator (S-IVM-1000, Scientifica, Uckfield, UK). After recording neuronal responses to all chosen sound stimulation protocols, the electrodes were slowly advanced deeper into the brain tissue to reach new recording positions. A single brain penetration was performed in each experiment. On average, Ins/S2 auditory responsive units were most frequent between 3.4 and 3.8 mm below the brain surface. At the end of the experiment, the end positions of electrodes in the brain were preserved during tissue fixation, making it possible to perform a precise histological reconstruction of the electrode tracks. For that purpose, the small microdrive with the electrode array was fixed to the skull with acrylic dental cement (Coral-fix, Tel Aviv, Israel). Once the cement cured, the microdrive was released from the micromanipulator. The rat was transferred to a fume hood for transcardial perfusion with the electrodes remaining in the brain.

### Neuropixels probe insertion

The 0.3-0.5 mm long slot in the dura was gently resected. The Neuropixels probe (version 3A) was slowly (less than 10 µm/s) inserted into the brain tissue while avoiding visible blood vessels to minimize trauma. A single-axis motorized micromanipulator (S-IVM-1000, Scientifica, Uckfield, UK) was used for insertion. Multiple electrode insertions (2-6) were made around the target coordinates. Neuronal responses to sounds were recorded with “bank 0” of the electrode - the 384 most distal electrodes (arranged over about 3.8 mm of the shank). The electrode targeting Ins/S2 was inserted approximately 4.2 to 5.1 mm below the brain surface. Broad-band noise and high-level tones (70-80 dB SPL) were used to determine the presence of auditory responses. Whenever auditory responsive units were found, the full set of sound stimulation protocols was presented. Protocols were recorded in: 23 out of 30 penetrations in Ins; 10 of 16 penetrations in A1/AC; 8 of 9 penetrations in MGB; 8 of 9 penetrations in IC. At the end of the experiment, the Neuropixels probe was removed from the brain tissue and the rat was transferred to a fume hood for transcardial perfusion.

### In vivo electrophysiology

Tungsten electrode recordings were performed using the AlphaLab SnR™ recording system (Alpha Omega Engineering, Nazareth, Israel). The raw voltage traces, captured at a sampling rate of 22 kHz, were then processed offline using Matlab (The Mathworks, Inc.). Neuropixels silicon probes were connected to a dedicated recording system (Phase 3A) as outlined on the website: https://github.com/cortex-lab/neuropixels/wiki/About_Neuropixels. Filtered and pre-processed voltage traces for action potentials (sampled at 30 kHz, filtered with a high-pass filter set at 300 Hz) and local field potentials (sampled at 2.5 kHz) were recorded, and subsequently underwent further offline analysis using Matlab.

### Auditory stimulation

The recordings were conducted in a soundproof chamber (IAC, Winchester, UK). Audio stimuli were generated using a custom-written Matlab (The Mathworks, Inc.). Sound signals were converted to voltage signals at a sampling rate of 192 kHz by a sound card (RME HDSP 9632, Haimhausen, Germany), attenuated (using a PA5 programmable attenuator, Tucker Davis Technologies - TDT, Alachua, FL, USA), and played through a two-channel electrostatic speaker driver (ED1, TDT) and an electrostatic speaker with coupler (EC1, from Tucker Davis Technologies). The speaker delivered sound through a flexible PVC tube placed in the right ear canal. The system was calibrated using a Knowles miniature microphone (EK3133-000, manufactured by Knowles, and calibrated relative to a B&K ¼” microphone) in the ears of several test rats. For pure tones, the attenuation level of 0 dB corresponded to 100 ± 10 dB SPL in the frequency range 1-30 kHz.

### Sounds and stimulation protocols

#### Broad-band noise

Responses to broad-band noise (BBN) bursts were collected by using a sequence of 280 BBN bursts, each with a duration of 200 ms, 10 ms linear onset and offset ramps, an inter-stimulus interval (ISI, onset-to-onset) of 500 ms, and seven different attenuation levels (0-60 dB at 10 dB steps). The attenuation levels were presented in a pseudo-random manner, so that each level was presented 40 times.

#### Frequency response

Frequency response areas (FRA) were determined by presenting quasi-random frequency sequences of 370 pure-tone bursts (with a duration of 50 ms, a 5 ms rise/fall time, and an ISI of 500 ms) at 37 frequencies (ranging from 1 to 64 kHz, with logarithmically spaced frequencies, 6/octave, and ten repeats each). The 370 pure-tone bursts were first presented at 0 dB attenuation, then again at attenuation levels that decreased by 10 dB steps until the threshold of neural activity was reached (usually at 50-70 dB attenuation). These data were used to determine the basic properties of auditory responses and to select a test frequency (TF) - a frequency that gave rise to the most consistent multi-unit responses at all sound levels. If activity was recorded from several neurons simultaneously, the minimum response threshold and TF were selected to match either the neuron with the most pronounced responses to pure tones or the most common TF and minimum response threshold among all recorded units. During recordings with Neuropixels silicon probes, one to three TFs were chosen per penetration based on the most prominent responses, and the rest of the protocol was presented multiple times, once for each TF.

#### Pure tone sequences

All sequences consisted of 500 tone presentations, with an inter-stimulus interval (ISI) of 300 ms. Each tone had a duration of 30 ms with 5 ms linear onset and offset ramps. The tones were presented at a constant level (usually ∼20 dB above the minimum threshold at the TF). Frequencies f1 and f2 (at a frequency separation Δf = *f*2/*f*1 = 1.44) were centered at the TF. However, since the different units recorded simultaneously could have different frequency tuning, for many units, f1 and f2 were not centered on their best frequency.

Seven test sequences that included frequencies f1 and f2 were used to study the habituation profiles of insular cortex units.

- In one oddball sequence, f1 occurred 475 times (95%, standard) and f2 25 times (5%, deviant); in the other, the roles of f1 and f2 were reversed.
- In the rare sequences, 475 trials (95%) consisted of silence only, and 25 trials (5%) had tones of frequency f1 or f2, respectively.
- The ‘Equal’ sequence contained equal numbers of f1 and f2 (50%, 250 presentations each).
- Two multi-tone sequences were recorded, called ‘diverse broad’ (12 equally spaced frequencies that included f1 and f2) and ‘diverse narrow’ (20 equally spaced frequencies covering a total range of about 1 octave).

The diverse broad sequence contained tones of frequencies f1 and f2 with a probability of 5% (25 times each). The other 450 stimuli of the sequence consisted of 10 different frequencies (presented 45 times each, with a 9% probability). These additional 10 tone frequencies were distributed below f1 and above f2, with successive frequencies separated by the same interval as f1 and f2 (frequency ratio of 1.44). The use of only 12 frequencies was necessary to avoid exceeding the frequency range of 1-64 kHz. To the extent possible, frequencies f1 and f2 were positioned in the middle of the frequency range used. Their position in the sequence was shifted when necessary to ensure that the overall range of frequencies tested did not exceed 1-64 kHz. In the diverse narrow sequence, tones of frequencies f1 and f2 were played together with 18 other tones, 25 times each. The 20 tones had logarithmically spaced frequencies with a ratio between the lowest and the highest tone in the sequence being 2.16 (slightly more than twice the distance between f1 and f2). These values were selected so that f1 was the 6th and f2 the 15th frequency in this sequence.

### Histological analyses

Unconscious animals received a lethal injection of sodium pentobarbital (900 mg i.p. per rat, Pentobarbital sodium 200 mg/ml, CTS Chemical Industries Ltd., Kiryat Malachi, Israel). When the rats stopped breathing, they were transcardially perfused in a fume hood with 350 ml of 0.1 M phosphate-buffered saline (PBS, Sigma Aldrich Inc., St. Louis, MO, USA) at room temperature, followed by 400 ml of 7% formaldehyde in 0.1 M PBS at around 4°C (Formaldehyde 35% w/w, Bio-Lab Ltd., Jerusalem, Israel). After perfusion, the brains were removed from the skulls and placed in 7% formaldehyde in 0.1 M PBS for at least 72 h at around 4°C. In rats implanted with tungsten electrode arrays, before removing the brain from the skull, the electrodes were slowly withdrawn from fixed brain tissue using the small microdrive still attached to the skull. Following fixation in formaldehyde, the brains were transferred to 30% sucrose, 4% formaldehyde solution in 0.1 M PBS at around 4°C (Sigma Aldrich Inc., St. Louis, MO, USA) for cryoprotection for about 7-14 days. The brains were levelled, and a small triangular longitudinal cut on the right hemisphere cortex was made (hemisphere marker). Each brain was placed in a polyethylene embedding mold (Peel-A-Way, Sigma Aldrich Inc., St. Louis, MO, USA). The mold was filled with optimal cutting temperature compound (OCT compound, Tissue-Tek, Alphen aan den Rijn, Netherlands). The polyethylene mold with the brain and OCT was placed in a custom aluminum freezing mold filled with a small amount of absolute ethanol (J.T.Baker, Deventer, Netherlands). The aluminum was then filled with dry ice to ensure that the brain froze rapidly. Frozen blocks with the levelled brains were transferred to a cryostat (CM1950, Leica). Consecutive 35 µm thick coronal sections were cut around the implantation site (left hemisphere). Brain slices were stained with green fluorescent Nissl stain (NeuroTrace® 500/525 Green Fluorescent Nissl Stain, Molecular Probes™, Eugene, OR, USA) according to the manufacturer’s instructions. The slices were then mounted onto slides (3 slices per slide) and after drying, covered with a mounting medium (Vectashield H1200, Vector Laboratories, Inc., Burlingame, CA, USA) and a cover glass. The slides were examined under standard fluorescent and bright field microscopy with 4x and 10x objectives. Multi-image array pictures were taken and arranged in a series of all consecutive sections around the implantation sites, from the most anterior to the most posterior. The positions of the electrode tips were estimated on the most relevant histological sections, and recording positions were proportionally derived.

### Quantification and statistical analysis

#### Preprocessing

22 sessions of Neuropixels recordings were analyzed for this study. Each session was initially sorted using Kilosort^75^ and then manually curated using phy (https://github.com/kwikteam/phy). In 11/23 penetrations, the non-curated units were first analyzed to look for auditory responsive units. These were used to determine the extent along the shank in which auditory activity was present, and only units in that region were manually curated.

#### Neuropixels-recorded units allocation

To locate the Neuropixels-recorded units within the brain, a 3^rd^ order polynomial was fitted to the shank trajectory on images of the histological sections, and units were located on the histological sections based on their relative location along the shank. In order to standardize the locations of the Neuropixels-recorded units along the insular-parietal field, we performed a linear transformation of the channel coordinates on the shank. This transformation set the point on the shank corresponding to the border between DI and GI as 0, and the border between GI and S2+ as 1. These linearly normalized distance values along the shank are termed “arc-distance” in the manuscript and allow us to compare unit locations consistently across different recordings.

#### Inhomogenous Poisson significant response test

For inclusion in the dataset, units were required to have a minimal firing rate (30 spikes in total for the BBN and FS protocols and 5 spikes in the 25 trials of the rare condition in the SSA protocol; spikes were counted over an interval from 100 ms before until 100 ms after the stimulus). Response significance was determined by a Poisson likelihood ratio test between a homogenous Poisson process whose rate was equal to the average activity during the 100 ms preceding stimulus onset, and an inhomogeneous Poisson process with rates that were computed in 10 windows of 10 ms each, for a total of 100 ms following stimulus onset. The logarithms of the probability ratios were summed to form a log likelihood ratio (LL). Under the null hypothesis, 2LL ∼ χ2 (10), and we required p<0.01 in this test.

#### Onset latencies

Trial-averaged responses were smoothed using a square filter of length 5 ms. We located the first local maximum following stimulus onset that was larger than the average spontaneous activity (100 ms before the onset of the stimulus) plus 7 times the standard deviation of the spontaneous activity’s as well as having a prominence (see https://www.mathworks.com/help/signal/ug/prominence.html) larger than 5% of the standard deviation of the spontaneous activity. We then located the local minimum preceding the local maximum, also requiring its prominence to be larger than 5% of the standard deviation in the spontaneous activity. Finally, we fitted a quadratic polynomial to the firing rates between these two, and interpolated the timepoint in which this quadratic function was equal to 25% of the local maximum. Finally, to correct for the “smearing” of the activity in time as a result of the smoothing filter, the estimated onset was shifted by 1.5 ms (30% of the 5 ms window used for smoothing). This timepoint was considered as the onset latency of the unit.

#### Frequency responses

To estimate the BF of the auditory responsive units from the FS stimulation protocol, we used a modification of the approach of Guo et al. (2012)^30^. The frequency that elicited the largest response in a 100 ms window post-stimulus, for the 3 lowest attenuations, was used as the BF.

#### SSA analyzed units

Units selected for the SSA analysis had to show significant responses using the inhomogenous Poisson significance test to at least one of the frequencies used when in the rare condition. Afterward, responses for each frequency in each condition were based on their average activity in a time window of 40 ms post-stimulus. Overall, 666 unit responses were analyzed from Ins/S2 and 355 from A1.

#### Adaptation in Narrow Frequency Channels (ANFC) Modeling analysis

Units for this analysis were chosen as a subset of the units chosen for the SSA analysis, with the additional requirement that the average response in the deviant condition be larger than the average response in the standard condition. Overall, 444 unit responses were used for Ins/S2 and 282 for A1. The model is based on ^43^. This model predicts the response to frequency *f*_0_, by estimating the resources used in a narrowly tuned frequency channel around *f*_0_. The model has two parameters: σ dictates the width of the frequency channels and τ is the time constant of the exponential resource recovery between stimulations. It can be shown ^43^ that given a fixed inter-stimulus interval *d* of a set of frequencies { *f* } played randomly with a respective set of probabilities { *p*_*f*_ }, the expected response to frequency *f*_0_ is:

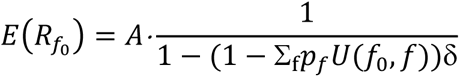

Where A is the ideal unadapted response to *f*_0_, *U*(*f*_0_, *f*) is the kernel that defines the adaptation channel centered on frequency *f*_0_, and 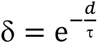. In our analysis, we used 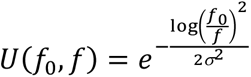. We normalized the responses to the rare responses, and therefore assumed that A=1. We then optimized the parameters *σ* & δ using MATLAB’s lsqnonlin function to approximate the responses in the other SSA conditions. The fit was computed for each unit separately.

## Declarations

### Author Contributions

Conceptualization, MMJ and IN; Experiments and histology, MMJ; Data analysis, MK; Data preprocessing, MK, MH, AP; Writing manuscript, MMJ, MK, IN; Supervision and funding acquisition, IN.

### Funding

This work was supported by AdERC grant GA-340063 (project RATLAND), by F.I.R.S.T. grant no. 1075/2013 from the Israel Science Foundation. IN holds the Milton and Brindell Gottlieb Chair in Brain Sciences.

### Ethics Approval and Consent to Participate

The experiments were carried out in accordance with the regulations of the ethics committee at The Hebrew University of Jerusalem (ethics approval no. NS-18-15652-4). The Hebrew University of Jerusalem is an Association for Assessment and Accreditation of Laboratory Animal Care (AAALAC) accredited institution.

## Acknowledgments

We thank Dr. Rony Kalman and Dr. Tamar Ravins and the staff of the Authority for Biological and Biomedical Models at The Hebrew University of Jerusalem for their excellent assistance in assuring high standards of animal welfare.

## Data Availability Statement

The data will be provided upon request.

## Competing Interests

The authors declare that they have no competing interests.

## Informed Consent Statement

Not applicable.

## Supplementary table

**Table S1.**
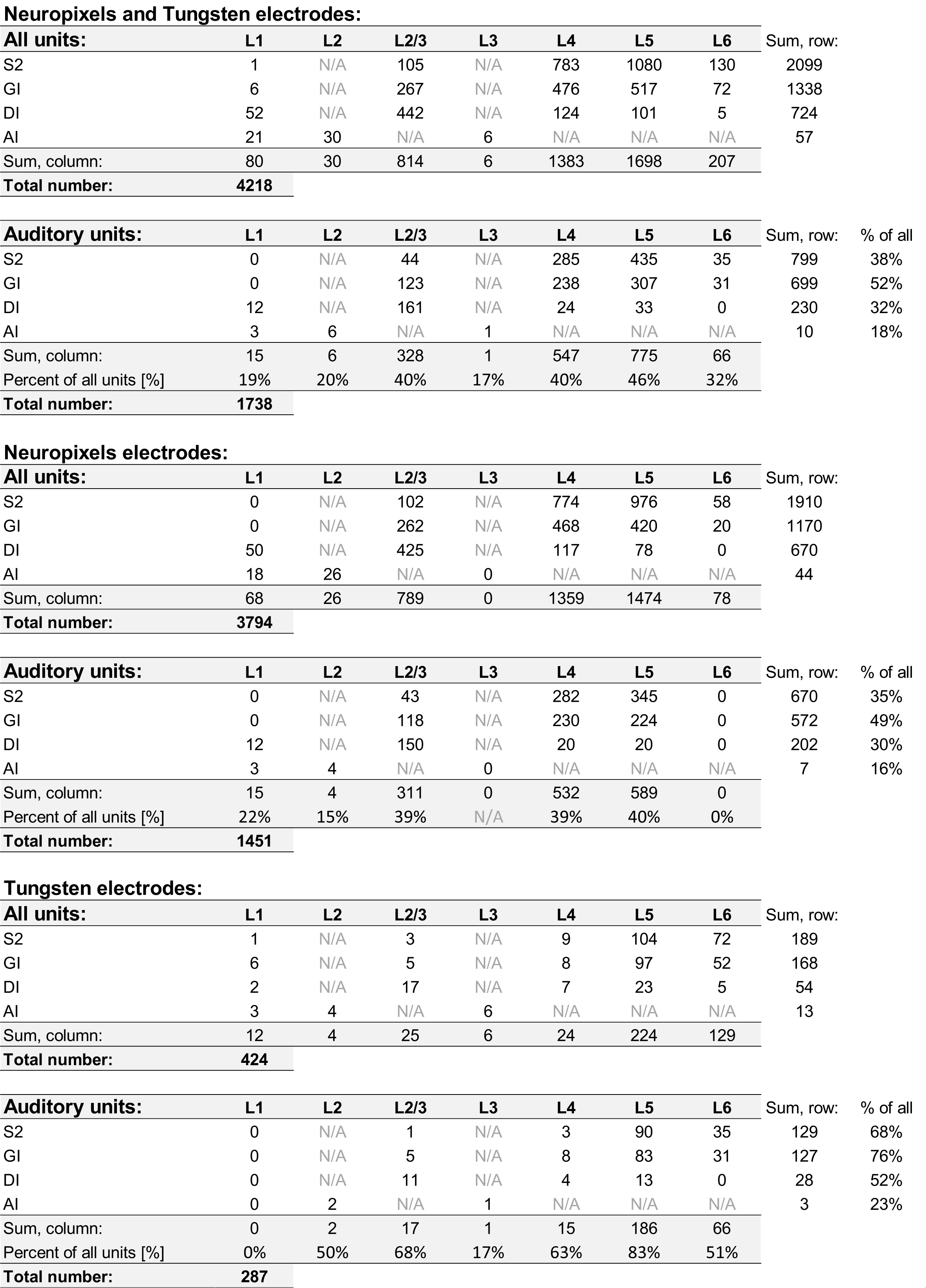
The numbers of all detected units and auditory cells, as well as their percentage among all units across cortical layers, in the secondary somatosensory cortex (S2) and granular insular cortex (GI), dysgranular insular cortex (DI), and agranular insular cortex (AI) were obtained using both Neuropixels and Tungsten electrodes.

## Notes

### Competing Interest Statement

The authors have declared no competing interest.

### Summary of Updates

Fig. 2 was changed for an updated version.

